# MetaVision3D: Automated Framework for the Generation of Spatial Metabolome Atlas in 3D

**DOI:** 10.1101/2023.11.27.568931

**Authors:** Xin Ma, Cameron J. Shedlock, Terrymar Medina, Roberto A. Ribas, Harrison A. Clarke, Tara R. Hawkinson, Praveen K. Dande, Lei Wu, Sara N. Burke, Matthew E. Merritt, Craig W. Vander Kooi, Matthew S. Gentry, Nirbhay N. Yadav, Li Chen, Ramon C. Sun

## Abstract

High-resolution spatial imaging is transforming our understanding of foundational biology. Spatial metabolomics is an emerging field that enables the dissection of the complex metabolic landscape and heterogeneity from a thin tissue section. Currently, spatial metabolism highlights the remarkable complexity in two-dimensional space and is poised to be extended into the three-dimensional world of biology. Here, we introduce MetaVision3D, a novel pipeline driven by computer vision techniques for the transformation of serial 2D MALDI mass spectrometry imaging sections into a high-resolution 3D spatial metabolome. Our framework employs advanced algorithms for image registration, normalization, and interpolation to enable the integration of serial 2D tissue sections, thereby generating a comprehensive 3D model of unique diverse metabolites across host tissues at mesoscale. As a proof of principle, MetaVision3D was utilized to generate the mouse brain 3D metabolome atlas (available at https://metavision3d.rc.ufl.edu/) as an interactive online database and web server to further advance brain metabolism and related research.

## Introduction

Spatial biology has emerged as a pivotal discipline for decoding the spatial organization and interactions of biomolecules within cells and tissues. Building upon the foundational pillars of spatial transcriptomics^1,2^, spatial proteomics^3,4^, and spatial metabolomics^5,6^, the field is significantly advancing our understanding of biological systems within multicellular organisms. Specifically, spatial metabolomics, performed through matrix-assisted laser desorption/ionization (MALDI) mass spectrometry imaging (MSI), enables the high-resolution mapping of metabolites in tissues at the mesoscale^6^. This methodology allows for the identification, quantification, and spatial distribution of multiple classes of metabolites such small molecules^7,8^, lipids^9,10^, glycogen and glycan-related complex carbohydrates^11,12^; thereby providing critical insights into cellular metabolism and disease mechanisms. Computational workflows are being developed to align and co-analyze high dimensional datasets produced from spatial metabolomics datasets^5,13,14^, bringing pathway enrichment and network analyses into spatial metabolism; and allowing for future advancements that promise a more comprehensive and nuanced understanding of cellular and tissue architecture. The field is thus poised for significant contributions to elucidating disease mechanisms and potential therapeutic interventions. For example, spatial metabolomics has provided insights into new skeletal myofiber subtypes^15^, dopaminergic lipid homeostasis during Parkinson’s disease^16^, and metabolic dependencies of glycogen during Ewing sarcoma^11^ and pulmonary fibrosis^17^.

Biological organisms and tissues operate in three-dimensional space. While a number of new technologies are presented for spatial transcriptomics^18-20^, current methodologies and data generated for spatial metabolomics are predominantly confined to two-dimensional analyses. The usefulness of 3D metabolomics has been demonstrated through manual co-registration and alignment of brightfield images^21-23^, but substantial manual input remains a major barrier for widespread adaptation and application of 3D metabolism data into an integrated workflow^24^. This limitation stands as a barrier to understanding the biological function of metabolic networks in 3D. One existing approach to 3D metabolic imaging is through magnetic resonance spectroscopy (MRS); however, this technique is constrained by a limited set of detectable metabolites and offers insufficient spatial resolution^25^. This is especially problematic in heterogeneous structures like the rodent brain in which the cellular composition of distinct brain regions changes at a spatial gradient that is significantly finer than the resolution offered by MRS. Advancing to a 3D metabolome at the mesoscale represents the next major milestone, serving to bridge the gap between 2D and 3D. Achieving 3D metabolomics at this scale would enable mapping of metabolic interactions and networks, reflecting the spatial complexities inherent in biological systems and bridging the gap in understanding from cellular to organismal levels. Such a shift from 2D to 3D would provide unprecedented insights into the spatial heterogeneity of metabolic activities within cells and tissues, facilitating a deeper understanding of the connection between molecules and physiology.

Here, we introduce MetaVision3D, a novel computational framework for the automated generation of 3D metabolome models from serial tissue sections. MetaVision3D represents a major advancement in spatial metabolomics, offering novel algorithms designed to reconstruct the three-dimensional metabolome of complex diseases or tissues. MetaVision3D employs a novel automated alignment framework through computer vision techniques (MetaAlign3D), normalization steps to account for intra-slice signal variabilities (MetaNorm3D), and then performs imputation (MetaImpute3D) and interpolation algorithms (MetaInterp3D) to improve overall 3D rendering visualization. The innovation lies both in the precision and resolution of the spatial data, and also in the workflow’s ability to compensate for experimental and technical variabilities inherent to MALDI imaging. MetaVision3D serves as a critical bridge between existing imaging modalities, enabling mesoscale spatial resolution and a comprehensive characterization of metabolic distributions within the brain. Leveraging MetaVision3D, we created the 3D mouse brain metabolome atlas in an interactive online database and web server (https://metavision3d.rc.ufl.edu).

## Results

### Alignment of 2D MALDI imaging datasets from serial tissue sections

The first step towards building a 3D spatial metabolomics atlas of the brain is the ability to seamlessly align serial tissue sections subjected to MALDI imaging in an automated fashion. Here, we developed a novel and powerful MALDI imaging alignment module (MetaAlign3D) in MetaVision3D based on the principle of enhanced correlation coefficient (ECC) maximization^26^, which has been adopted for image alignment in the field of computer vision. Using this algorithm, MALDI images are systematically aligned through an iterative process that maximizes the correlation coefficient between adjacent sections directly from data matrix of MALDI images. Utilizing the first section as the starting point, each subsequent image was subjected to parametric transformations—comprising rotations, translations, and in some instances, scaling and skewing—to identify the optimal overlay (Fig. 1A). The ECC maximization principle functioned as the guiding metric in this process, providing a quantitative measure of similarity that was optimized until the alignment of molecular patterns across serial sections reached the peak of the correlation function (see methods). To test MetaAlign3D, we applied it to align in five sequential sagittal-cut MALDI image sections from the medial plan of a mouse hemi brain (Fig. 1B and supplemental Fig. 1A). Comparing to manual fit (see methods), MetaAlign3D achieved superior alignment (Fig. 1B, supplemental Fig .1A). This was evidenced by the enhanced overlap accuracy of the anatomical landmarks highlighted by the metabolite distribution across the brain section (Fig 1B and supplemental Video 1).

**Figure 1.**
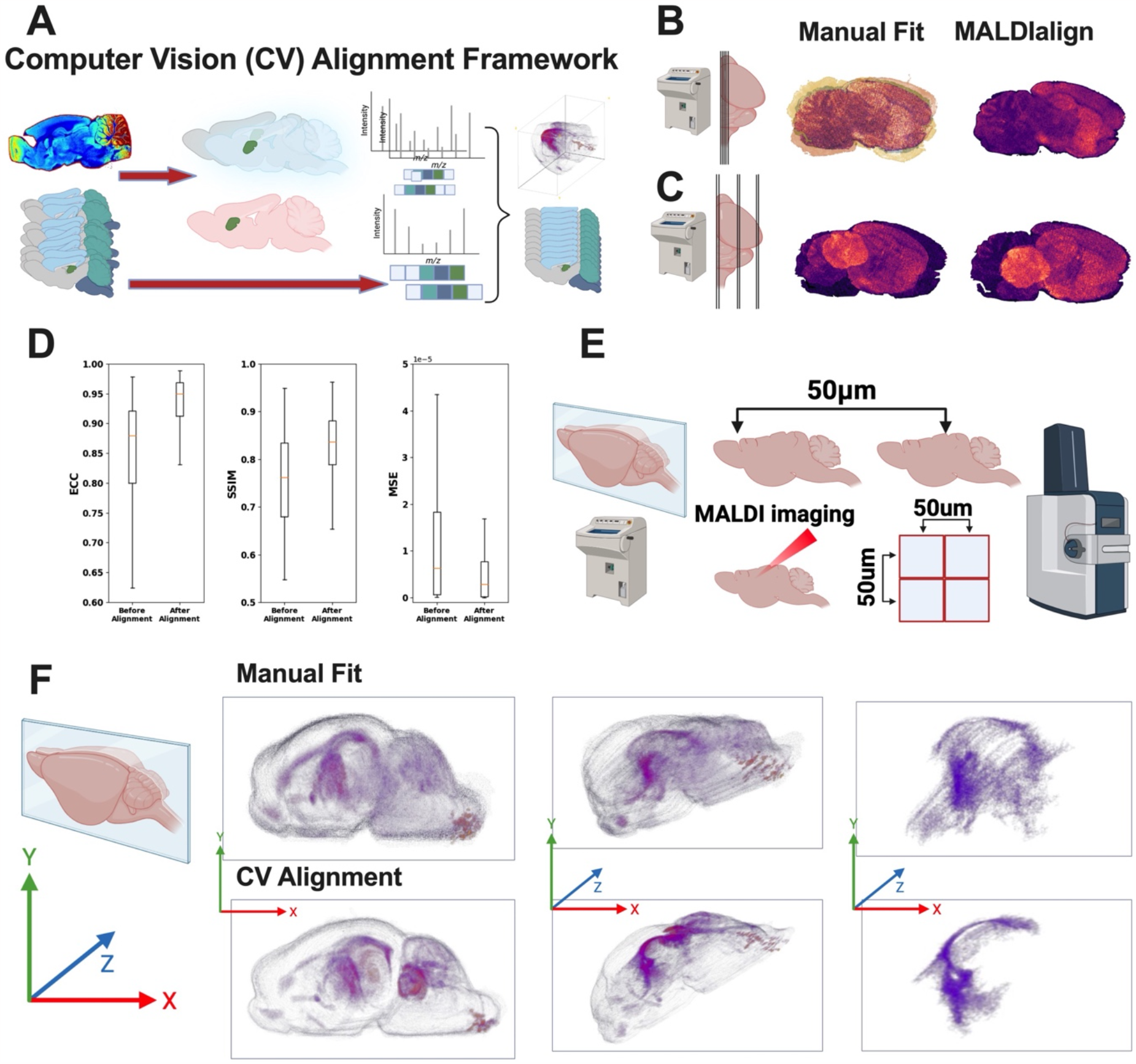
MALDIalign for the automated alignment of serial MALDI imaging tissue sections. **a** Schematics computation workflow for the automated alignment of sequential MALDI imaging tissue powered by computer vision. **b** MALDIalign vs. manual fit for five serial sagittal sections cut from the medial side of a mouse hemi brain. **c**. MALDIalign vs. manual fit for six sagittal sections cut from the medial side to lateral side of a mouse hemi brain with major shift in tissue size. **d**. Statistical measures of alignment and fit quality between manual fit and MALDIalign, enhanced correlation coefficient (ECC), structural similarity index measure (SSIM), and mean squared error (MSE). **e**. Schematics experimental workflow MALDI imaging of mouse hemi brain at 50um resolution. total of 79 sections were acquired. **f**. MALDIalign vs. manual fit for the total 79 serial sagittal sections cut from a mouse hemi brain for the creation of spatial metabolome Atlas in 3D.

To test the ability of MetaAlign3D to fit serial tissue sections of different sizes, we performed MALDI imaging on tissue sections derived from the medial and lateral side of the mouse sagittal hemisected brain (Fig. 1C and supplemental Fig. 1B). MetaAlign3D corrected for distortion due to size differences between lateral and medial brain sections and maintained anatomical alignment highlighted by metabolic features (Fig. 1C). This improvement is particularly evident when juxtaposed against manual alignment efforts, where notable imperfections were observed (Fig. 1C). Statistical measures of alignment and fit quality, including enhanced correlation coefficient (ECC) for geometric transformation^26^, structural similarity index measure (SSIM) were used to compare image similarities^27^, and mean squared error (MSE) that quantified differences between serial images^28^. MetaAlign3D of the entire volume demonstrates substantial improvement over the manual fitting technique for all parameters assessed (Fig. 1D).

### Construction of the 3D spatial metabolomics atlas of the brain

We further demonstrate the scope and utility of MetaAlign3D in an ambitious task of constructing the first mesoscale mouse brain 3D metabolomics atlas. To achieve this, we performed cryosectioning of 10 μm thick brain sections from a mouse hemi-brain. Importantly, sections are 50 μm apart (z-axis), corresponding to the MALDI imaging spatial resolution of 50 μm (x- and y-axis). This strategy allows creation of a 3D brain atlas at 50 μm resolution (Fig. 1E). A total of 79 serial brain sections were collected from cryosectioning and subjected to MALDI imaging. We employed MetaAlign3D and manual fit to connect all 79 brain sections and created a 3D rendering of the metabolic features (see method). We rendered LPE 16:1 in 3D view using ImageJ (Fig. 1F, see methods) to allow visualization of detailed brain anatomical structures (Fig. 1F). For example, the structure of corpus collosum is much more pronounced after MetaAlign3D compared to manual fit (Fig. 1F and supplemental Video 1).

### Normalization, imputation, interpolation to improve visualization of the 3D atlas

Upon completion of the alignment, a series of challenges became apparent. First, due to the inherent characteristics of the MALDI workflow, discernable variabilities were observed in certain metabolic features both within and between slides even after normalization to total ion current (TIC) (supplemental Fig. 2). Such variations pose challenges for ensuring consistent representations of the metabolic landscapes across sections. Additionally, the imperfections associated with handling and sectioning frozen brain tissue at 10 μm became evident, as several sections exhibited missing regions—likely attributed to the fragile nature of the frozen brain (supplemental Fig. 4A). These findings underscore the complexities involved in the experimental aspects of such atlas construction and point towards areas requiring further refinement and methodological adaptation.

While MALDI imaging analyses were normalized to TIC, inter-slide normalizations are not commonly performed. Inter-slide variability is often caused by the MALDI instrument, leading to inconsistencies in the representation of metabolites across serial sections (supplemental Fig. 2A). Such disparities could potentially skew the interpretative outcomes of the atlas. To account for these issues, we instituted a normalization strategy for each metabolite, using the median value of the section as the reference point (see methods). By normalization to the tissue section median, we markedly improved the inter-section consistency (supplemental Fig. 2B). This ensured a smoother transition of metabolite intensities between serial sections. As a result, the normalized dataset provided a more contiguous and coherent display of the metabolic landscape, substantially reducing the technical variabilities associated with the MALDI process. lipids PIP 38:4 and PI 36:4 are shown as examples (supplemental Fig. 3).

To address the challenges stemming from the physical handling of brain sections, which occasionally led to missing regions or gaps in the tissue, we developed an advanced imputation module named MetaImpute3D in MetaVision3D to fill in these gaps (supplemental Fig. 4A). For this process, we utilized two adjacent sections - one from either side of the compromised section - as references. Drawing data from these intact sections, a linear equation was employed to generate and create pixel values for the missing regions, creating a digital fill for the physical gaps (see methods). The efficacy of this imputation approach was striking; post-imputation, the once-damaged tissue sections appeared almost indistinguishable from their undamaged counterparts (supplemental Fig. 4C-D).

Building on this, we implemented computer vision to smoothly interpolate between sections. Similar to imputation, this was achieved by calculating the mean of corresponding pixels from four adjacent tissue sections (two on each side), effectively creating interpolated sections that bridge the gaps between the original slices (supplemental Fig. 5A-B). By implementing this interpolation, named MetaInterp3D in MetaVision3D, as an optional step in our analysis, we were able to introduce two additional sections between each original section, expanding our dataset to a total of 237 sections (supplemental Fig. 5C-D). This significantly enhanced z-axis continuity, thereby enabling optimal 3D reconstruction. It is worth noting that the addition of interpolated sections was methodologically sound, as statistical measures of alignment were evaluated to confirm that the process of interpolation did not introduce any artifacts or alignment errors. The comparative analysis post-interpolation maintained the high standards of alignment accuracy using SSIM, ECC, and MSE (supplemental Fig. 5E).

### 3D mouse brain metabolomic atlas as an online resource

We further produce the first 3D spatial metabolomics atlas of the mouse hemi-brain using MetaVision3D (Fig. 2A). MetaVision3D data output is compatible with ImageJ (see methods), a widely used image processing program, which researchers can utilize to visualize 3D models of individual metabolites (supplemental video 2-10). Through projection or volume viewing techniques available within ImageJ^29^, users can generate detailed three-dimensional visualizations that can be viewed and studied from various angles (supplemental Fig. 6A). For example, PE 44:12 is observed to be enriched in the molecular and granular layers of the cerebellum (Fig. 2B, supplemental videos 2-4), PS 38:6 is almost exclusively enrichment in the dentate gyrus (Fig. 2C, supplemental videos 5-7), similarly both PI 38:5 and PS 40:6 are enriched in the frontal cortex, and granular layers of the hippocampus and cerebellum (supplemental Fig. 6B-C and supplemental videos 8-10). These metabolites are visualized using both top-down (supplemental videos 3,6,9) and frontal views (supplemental videos 2,5,8), offering a comprehensive perspective that highlights spatial landmarks and the three-dimensional organization of metabolic compounds. Importantly, we made this atlas accessible through a dual-platform approach: as an interactive website portal (https://metavision3d.rc.ufl.edu) and as a downloadable database. The web-based platform offers a user-friendly interface where each metabolite is represented through both 2D slice transitions and 3D renderings with a customizable pixel intensity threshold (Fig. 2D). This interactive feature allows for an intuitive exploration of the metabolomic landscape, providing users with the ability to navigate through the brain’s structure in two dimensions, or to delve into the three-dimensional organization of metabolites, delivering insights into the intricate spatial relationships that exist within the brain.

**Figure 2.**
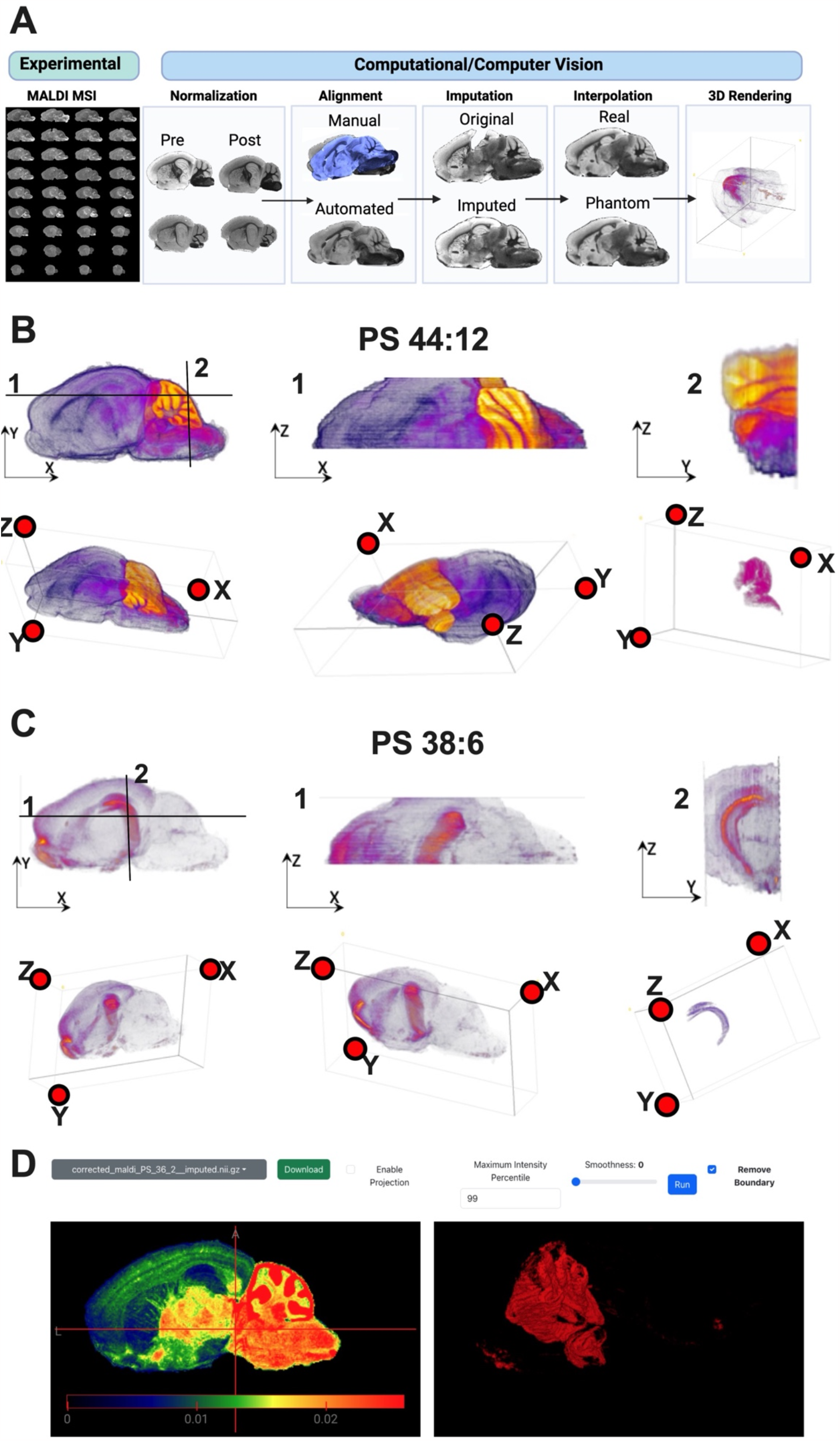
Generation of the spatial metabolome atlas of by MetaVision3D. **a**. Computational framework for MetaVision3D for the generation of spatial metabolome of mouse hemi brain in 3D. **b**. MetaVision3D rendering of PS 44:12 in 3D space. Top down and front cross section views are designated as 1 and 2 shown on the 2D side view image. 3D rendering is visualized by ImageJ using both projection mode and 3D fill of regions with high intensity of PS 44:12 in the cerebellum (see method). **c**. MetaVision3D rendering of PS 38:6 in 3D space. Top down and front cross section views are designated as 1 and 2 shown on the 2D side view image. 3D rendering is visualized by ImageJ using both projection mode and 3D fill of regions with high intensity of PS 38:6 in the cerebellum (see method). **d**. Screenshot of https://metavision3d.rc.ufl.edu/ and its functionalities.

## Discussion

In this study, we introduced MetaVision3D, a novel workflow designed to facilitate the generation of the 3D metabolome of tissues. MetaVision3D integrates a series of functionalities through computer vision—including alignment, imputation, and interpolation to construct spatial metabolome in 3D. Leveraging MetaVision3D, we constructed the first 3D metabolome of the mouse brain at mesoscale. The innovation lies both in the precision and resolution of the spatial data and also in the workflow’s ability to compensate for experimental and technical variabilities inherent to MALDI imaging. The shift from conventional 2D representations to a 3D atlas marks a significant technological leap, providing an unprecedented depth of metabolic insight that mirrors the complex three-dimensional structure of the brain. By moving into the third dimension, Metavision3D allows for an understanding of metabolic distributions within the brain’s spatial architecture, setting a new standard for spatial metabolomics and opening new avenues for neurological research at mesoscale. Finally, to facilitate brain metabolism research, we are publishing the mouse brain metabolome atlas generated from MetaVision3D through an interactive website as well as a downloadable database.

Building on the successful alignment of serial brain sections, MetaVision 3D demonstrates a detailed spatial characterization of metabolites that correlate with distinct neuroanatomical regions. For instance, the lipid species PE 44:12 was observed to be enriched within the molecular and granular layers of the cerebellum, implicating its possible role in the unique metabolic demands in motor and cognitive function^30^. Similarly, PS 38:6 displayed a striking pattern of enrichment almost exclusively in the dentate gyrus, suggesting a specialized function in this hippocampal subregion known for its involvement in memory formation and spatial navigation^31^. The distribution of phosphoinositides such as PI 38:5, along with PS 40:6, were predominantly localized to the frontal cortex, a region implicated in executive higher cognitive functions. These phospholipids also showed a notable presence in the granular layers of the hippocampus and cerebellum, areas critical for synaptic plasticity and motor control, respectively^32,33^. These spatially resolved metabolic profiles, revealed by MetaVision3D, underscore the inseparable relationship between metabolite distribution and brain functionality. The precise localization of these metabolites could open new avenues for investigating their specific roles in neural metabolism and the pathophysiology of neurological disorders and should be investigated in detail in the future.

Armed with this tool, there are a number of paths to pursue in the future. For example, expanding the database of our 3D metabolomic atlas to a wider range of molecular classes such as small molecule metabolites, glycogen-, and glycan-related complex carbohydrates. Another exciting revenue is to apply MetaVision3D to explore the 3D metabolism of other organs such as the liver, kidney, heart, and muscles would be informative for whole body physiology, and eventually in creating the human body atlas. Leveraging MetaVision3D to examine the 3D metabolic dysregulation in chronic and acute disease models and define how metabolism changes during aging and sex are critical for our understanding of metabolic etiology of diseases. In parallel, integration of metabolomic data with other omics layers in 3D, such as genomics, transcriptomics, and proteomics, that would create a multi-omics atlas that offers a more holistic view of the biological systems at play, this effort is already under way for 2D spatial modalities^34,35^. Ultimately, the convergence of these data streams through advanced computational analyses will lead to a deeper understanding of the molecular underpinnings of diseases and aid in the development of targeted therapeutic interventions. The 3D atlas is not just an endpoint but a foundational tool for the next era of biological research that connects our understandings of phenotype to physiology.

## ACKNOWLEDGMENTS

This study was supported by National Institute of Health (NIH) grants R01AG066653, R01CA266004, R01AG078702, RM1NS133593 to R.C.S., R35NS116824 to M.S.G., R35GM142701 to L.C

