## Supplementary material for "MetaVision3D: Automated Framework for the Generation of Spatial Metabolome Atlas in 3D": sup figures 1-6

[1] Department of Biochemistry & Molecular Biology, College of Medicine, University of Florida, Gainesville, FL, USA
[2] Center for Advanced Spatial Biomolecule Research, University of Florida, Gainesville, FL, USA
[3] Department of Biostatistics College of Public Health and Health Professions & College of Medicine, University of Florida, Gainesville, FL, USA
[4] Department of Neuroscience, University of Florida, Gainesville, FL, USA
[5] McKnight Brain Institute, University of Florida, Gainesville, FL, USA
[6] Center for Translational Research in Neurodegenerative Disease (CTRND), University of Florida, Gainesville, FL, USA
[7] Institute on Aging, University of Florida, Gainesville, FL, USA
[8]Kennedy Krieger Institute, Baltimore, MD, USA
[9]Johns Hopkins University, Baltimore, MD, USA
### These authors contributed equally: Xin Ma; Cameron J. Shedlock; Terrymar Medina; Roberto A. Ribas
* These authors jointly supervised this work: Li Chen; Ramon C. Sun

#### Supplementary Figure 1-6

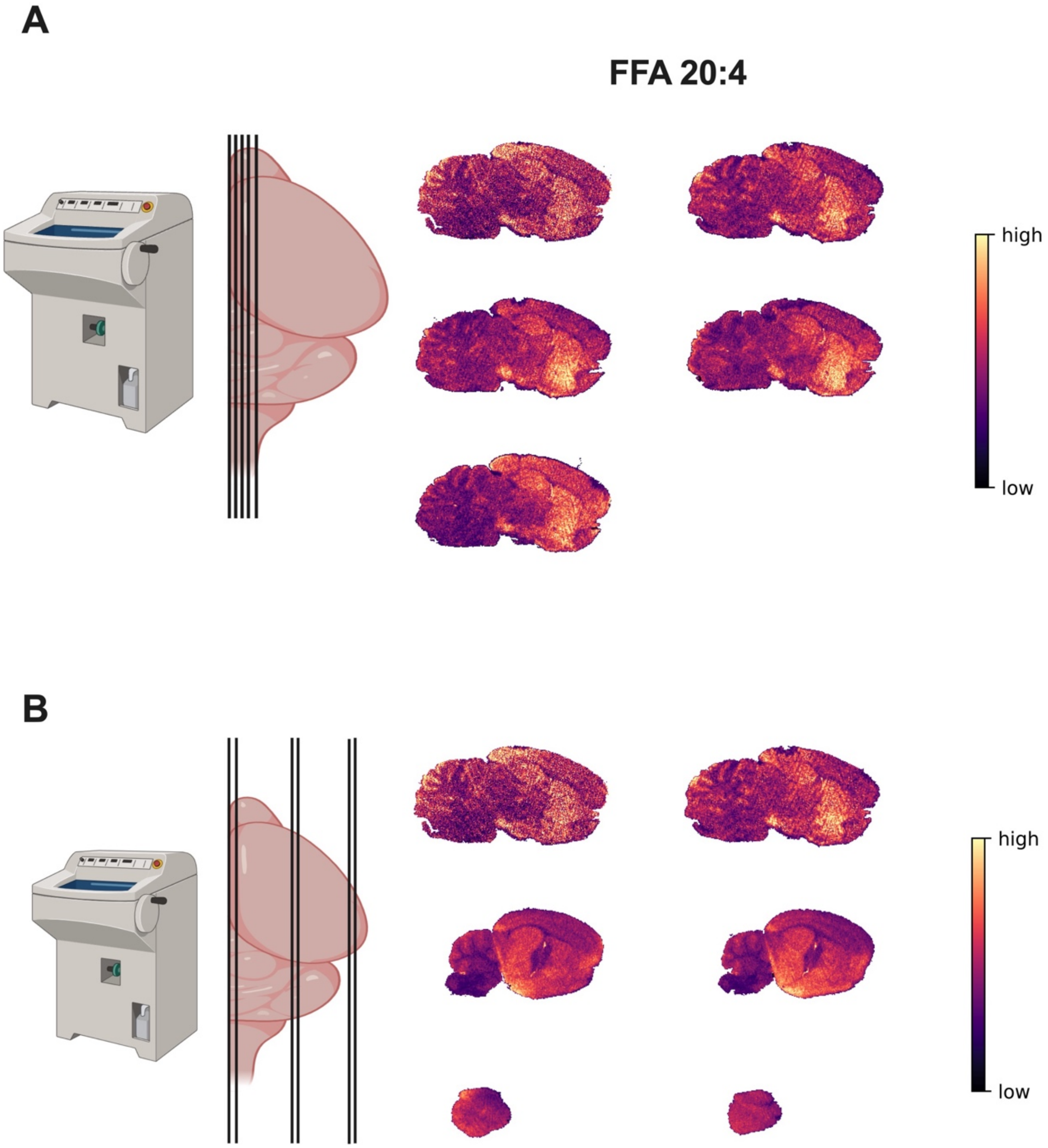


**Supplemental Figure 1. 2D MALDI heatmap images of serial mouse brain sagittal sections.**
**a.** Schematic of five serial sagittal cut brain sections from the medial plane (left). Heatmap images for the spatial distribution of the free fatty acid (FFA) 20:4 in 2D for all five serial brain sections tested for alignment. **b.** Schematic of serial sagittal cut brain sections from the medial to posterior brain with different section sizes (left). Heatmap images for the spatial distribution of the free fatty acid (FFA) 20:4 in 2D for all six serial brain sections with different sizes tested for alignment.

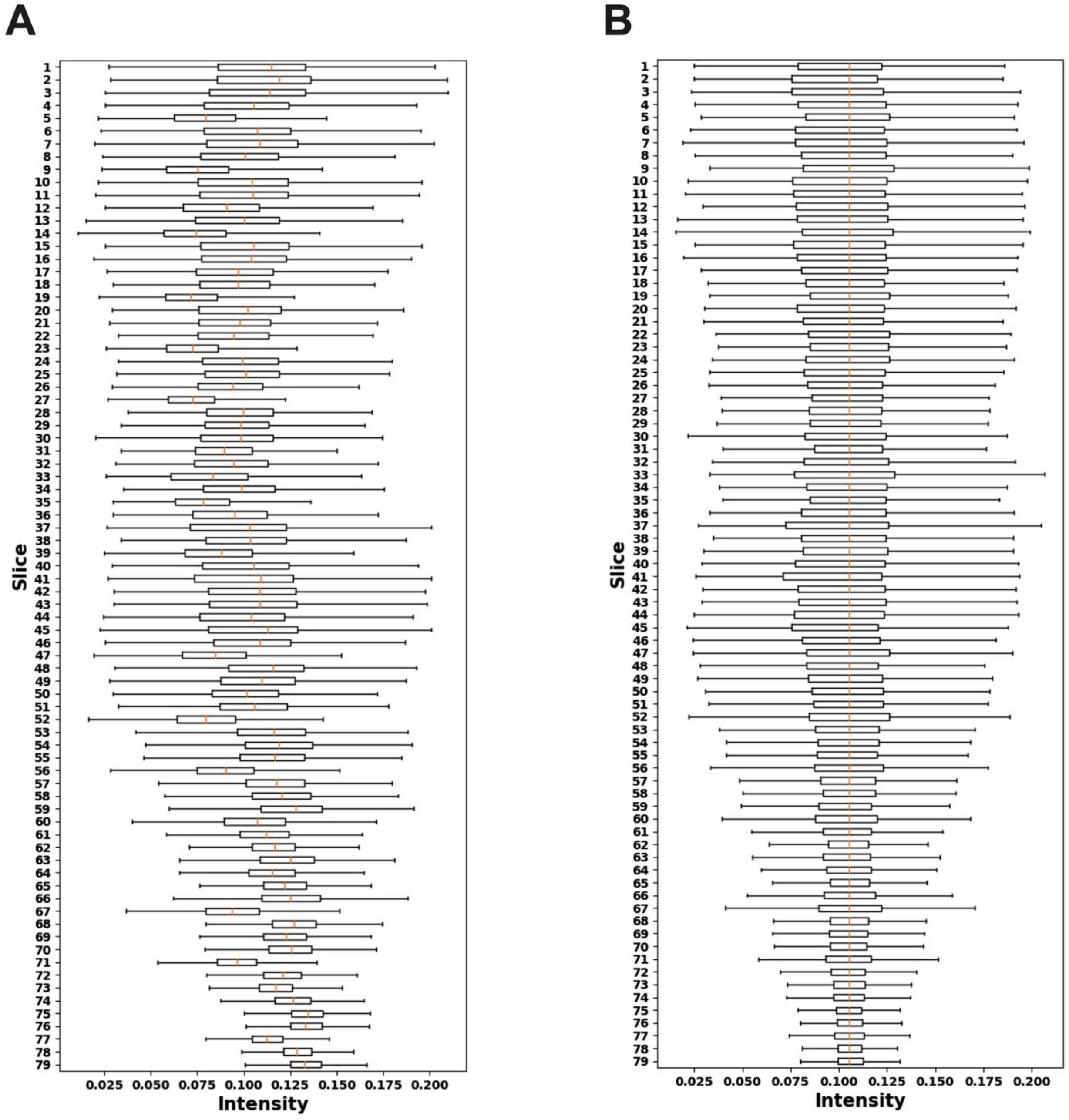


**Supplemental Figure 2. Normalization of serial brain sections for 3D construction using MetaNorm3D**

**a.** Representative intensity distribution of PIP 38:4 among all 79 serial sections of the mouse sagittal hemi-brain. **b.** Representative intensity distribution of PIP 38:4 after slide normalization (see method) among all 79 serial sections of the mouse sagittal hemi-brain.

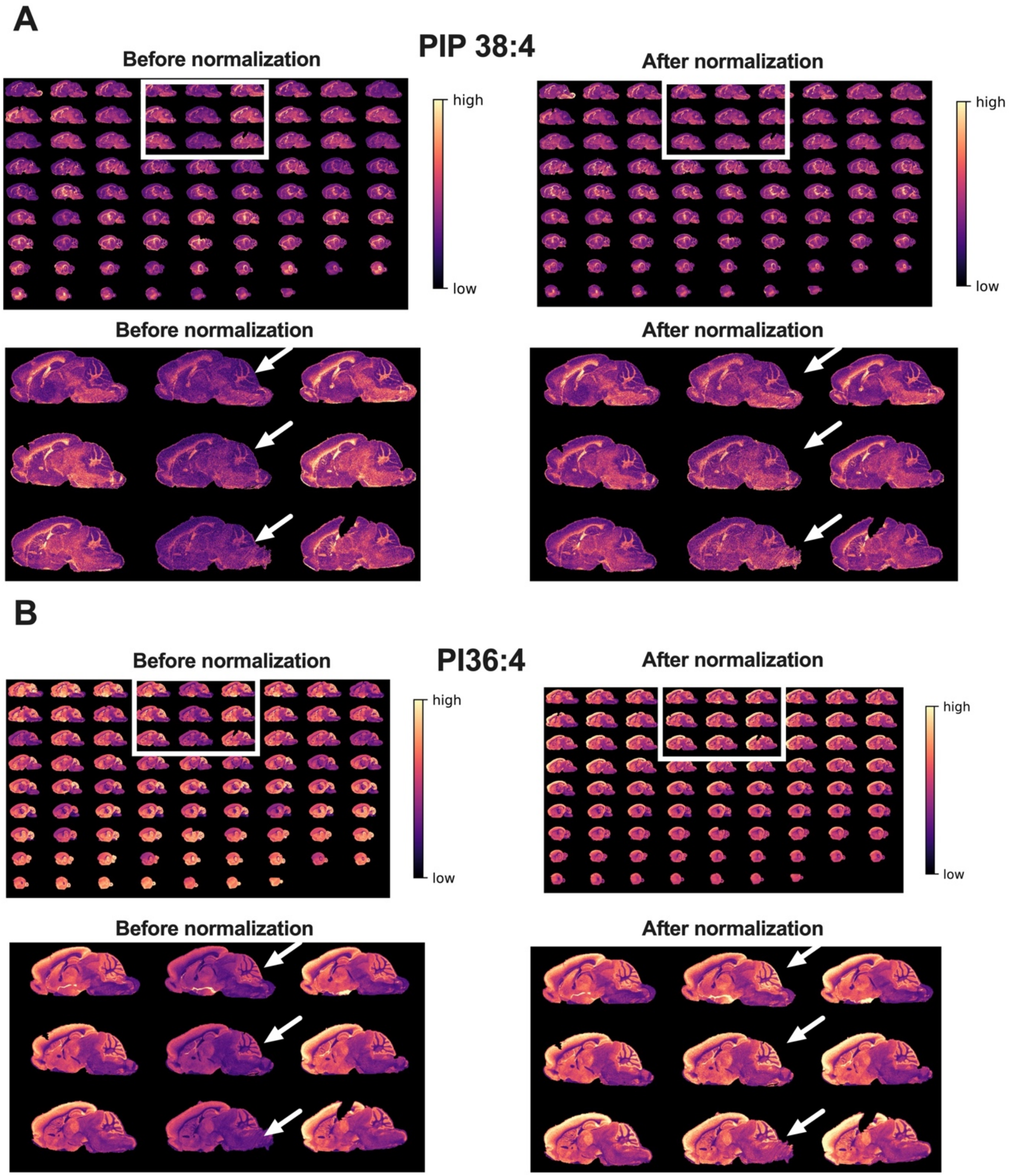


**Supplemental Figure 3. 2D MALDI heatmap images of serial mouse brain sagittal sections before and after normalization. a.** Heatmap images for the spatial distribution of PIP 38:4 in 2D for all 79 serial brain sections for 3D construction before and after normalization. Insert sections represents tissue sections with disparities in total abundance are zoomed in below. **b.** Heatmap images for the spatial distribution of PI 36:4 in 2D for all 79 serial brain sections for 3D construction before and after normalization. Insert sections represents tissue sections with disparities in total abundance are zoomed in below.

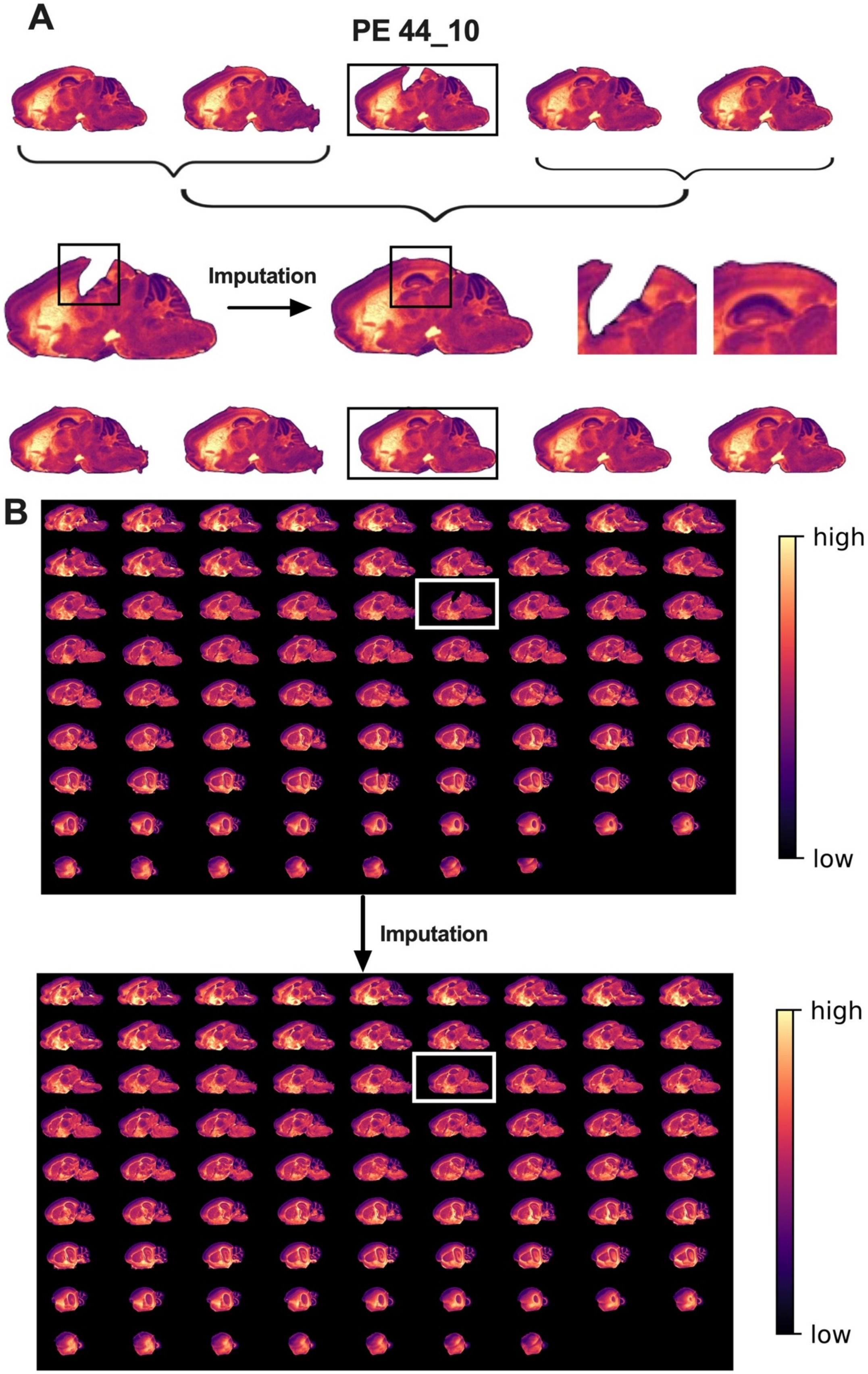

A
PE 44_10
Imputation
B
high
low
Imputation
high
low

**Supplemental Figure 4. MetaImpute3D to fill in imperfections during sample handling.**
**a.** Schematic of imputation pipeline by leverage anterior and posterior serial sections to fill in gaps in tissue from experimental imperfections. **b.** Heatmap images for the spatial distribution of PE 44:10 in 2D for all 79 serial brain sections for 3D after normalization, but before and after imputation of gaps in tissue. A representative tissue section after imputation is highlighted in the insert.

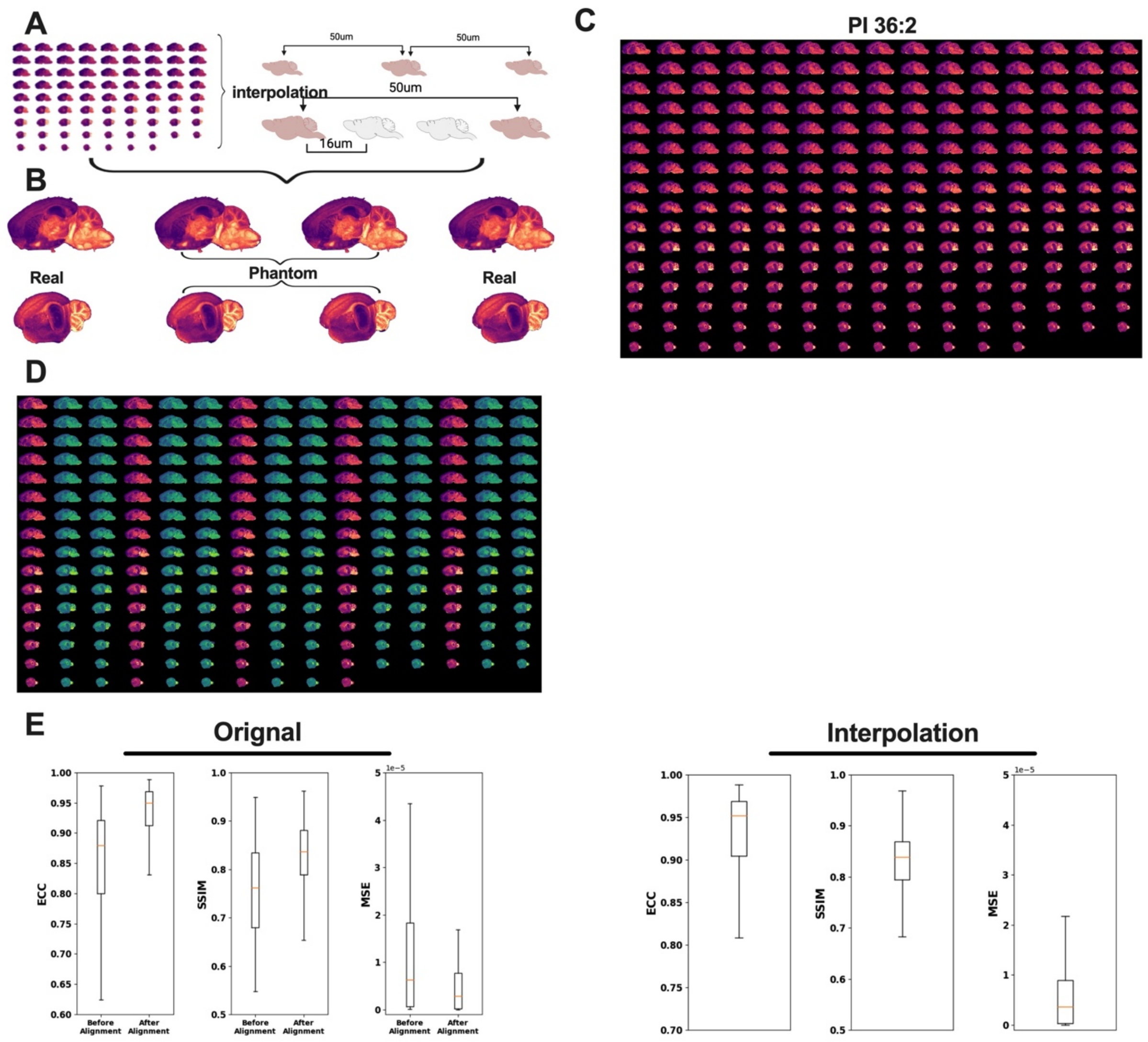


**Supplemental Figure 5. MetaInterp3D to fill in imperfections during sample handling. a.** Schematic of interpolation pipeline by leverage anterior and posterior serial sections to create phantom tissue sections to improve Z-axis resolution. **b.** Heatmap images for the spatial distribution of PEI 36:2 in 2D for 4 serial brain sections from medial and posterior brain regions. Phantom brain sections are labeled. **c.** Representatives heatmap images for the spatial distribution of PI 36:2 in 2D for all 79 real and 158 phantom serial brain sections for 3D construction. **d.** Representatives heatmap images for the spatial distribution of PI 36:2 in 2D for all 79 real and 158 phantom serial brain sections for 3D construction, phantom sections are highlighted in green. **e**. Statistical measures of alignment and fit quality measured by enhanced correlation coefficient (ECC), structural similarity index measure (SSIM), and mean squared error (MSE) before (79 sections) and after interpolation (316 section).

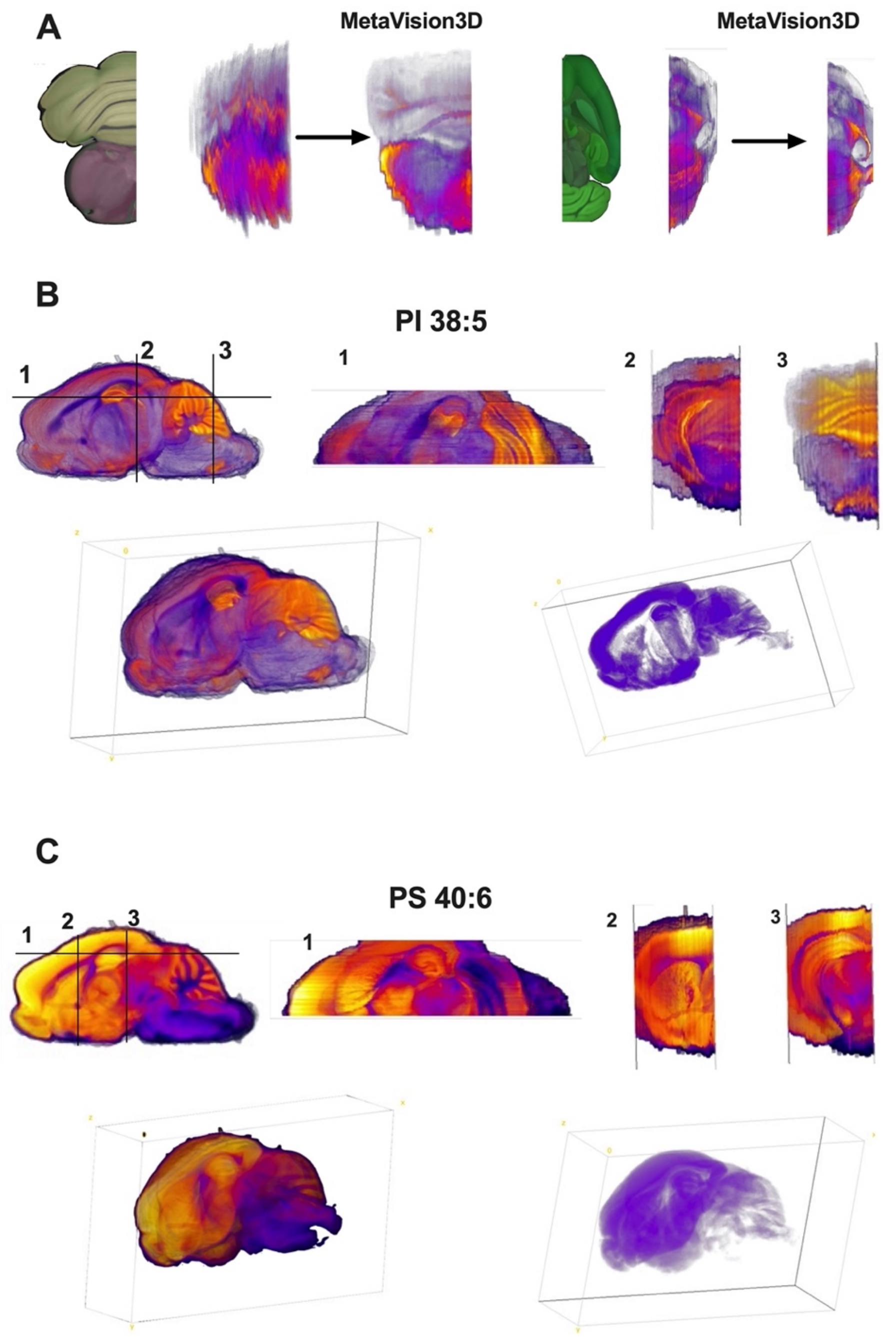


**Supplemental Figure 6. Additional examples of spatial metabolome atlas of by MetaVision3D.** **a**. Front cross section and top-down view of fine brain anatomical structures of mouse brain cerebellum and hippocampal region in manual fit and after MetaVision3D framework. **b and c**. MetaVision3D rendering of PI 38:5 and PS 40:6 in 3D space. Top down (transverse) and front cross section (coronal) views are designated as 1, 2 or 3 shown on the 2D side view image. 3D rendering is visualized by ImageJ using both projection mode and 3D fill of regions with high intensity of both features.
